# Visual Clustering of Transcriptomic Data from Primary and Metastatic Tumors – Dependencies and Novel Pitfalls

**DOI:** 10.1101/2021.12.03.471112

**Authors:** André Marquardt, Philip Kollmannsberger, Markus Krebs, Antonella Argentiero, Markus Knott, Antonio Giovanni Solimando, Alexander Kerscher

## Abstract

Personalized Oncology is a rapidly evolving area and offers cancer patients therapy options more specific than ever. Yet, there is still a lack of understanding regarding transcriptomic similarities or differences of metastases and corresponding primary sites. Applying two unsupervised dimension reduction methods (t-Distributed Stochastic Neighbor Embedding (t-SNE) and Uniform Manifold Approximation Projection (UMAP)) on three datasets of metastases (n=682 samples) with three different data transformations (unprocessed, log10 as well as log10+1 transformed values), we visualized potential underlying clusters. Additionally, we analyzed two datasets (n=616 samples) containing metastases and primary tumors of one entity, to point out potential familiarities. Using these methods, no tight link between site of resection and cluster formation outcome could be demonstrated, neither for datasets consisting of solely metastasis nor mixed datasets. Instead, dimension reduction methods and data transformation significantly impacted visual clustering results. Our findings strongly suggest data transformation to be considered as another key element in interpretation of visual clustering approaches along with initialization and different parameters. Furthermore, the results indicate only minor transcriptional differences for metastases and corresponding primary tumors.

## 1. Introduction

From a clinical perspective, characteristic metastatic patterns frequently occur for specific cancer entities [1]. Thus, site of metastasis has a considerable effect on patients’ prognosis. For example, liver metastases derived from pancreatic adenocarcinoma are prognostically worse than lymph node or lung metastases [2,3]. Still, it is unclear whether there is a biological or genetic determination for tumors to develop regional metastases or even distant metastasis with preferred target regions [4–6].

From a biological point of view, tumor cells develop through clonal evolution, favoring tumor heterogeneity reflected by different driver mutations or genomic alterations. Due to the accumulation of different alterations, metastases can occur. However, circumstances of local or distant metastases still need to be investigated, as it was already shown that distant metastasis can develop without previous local metastasis [7]. Additionally, transcriptomic differences of the linear [8] and the parallel progression [9] model of metastasis are still not fully clarified.

Due to the advances in personalized oncology, comprehensive elucidation of primary tumors and metastases is an ongoing process. Still, the determination of possible transcriptomic differences or similarities between metastases – especially from several different resection sites – and primary tumors is an unmet need. Previous approaches frequently analyzed the differences between two specific groups, e.g. bone and brain metastasis [10], or the mutational evolution, while showing a high concordance of primary and metastatic tumors [11–13]. As a result, a comprehensive study of the transcriptomic characteristics of metastases and corresponding primary tumors is lacking.

Visual clustering – based on data dimension reduction methods – is one potential approach to determine transcriptomic differences and proximities of different metastasis sites. The mainly used visualization methods for this purpose are t-Distributed Stochastic Neighbor Embedding (t-SNE) [14] and Uniform Manifold Approximation and Projection (UMAP) [15]. They have already been widely applied in the field of single cell sequencing [16–18] but also bulk RNA-sequencing [19–22] to visually separate transcriptionally similar cell populations from diverging populations in a two-dimensional space. Furthermore, recent studies have shown the critical impact of initialization [23] and parameters [24] on data dimension reduction methods.

To search for transcriptomic dependencies caused by the site of metastasis, t-SNE and UMAP were used to analyze three metastasis datasets – prostate cancer (PCa), neuroendocrine PCa, and skin cutaneous melanoma, totaling in 682 samples. For a comprehensive analysis, unprocessed Fragments Per Kilobase Million (FPKM) values, – obtained after normalization of the mapped sequencing reads, as well as log10 and log10+1 transformed data were analyzed, as logarithmic transformations are commonly used when analyzing gene expressions.

## 2. Materials and Methods

### 2.1. Data acquisition

RNA sequencing data from three different metastasis datasets were analyzed. The first dataset contained n=266 samples from metastatic prostate carcinoma (PRAD-SU2C - Dream Team [25]), the second consisted of n=49 samples from metastatic neuroendocrine prostate carcinoma (NEPC-WCM [26], and the third dataset consisted of n=367 metastatic skin cutaneous melanomas (TCGA-SKCM-Metastatic [27]). All datasets indicate the site of resection, which served as the basis for further analyses.

For evaluation purposes, we used an additional dataset known to form distinct clusters based on histopathological subgroups. The TCGA-KIPAN dataset, consisting of three renal cell carcinoma (RCC) subgroups – TCGA-KIRC (clear cell RCC, n=538), TCGA-KIRP (papillary RCC, n=288), and TCGA-KICH (chromophobe RCC, n=65).

For further testing similarities and differences between primary and metastatic tumors, we used the complete TCGA-SKCM dataset – adding n=103 primary tumor samples for a total of n=470 samples – and a metastatic breast cancer dataset (MBCproject; cBioPortal [28,29] data version February 2020 [30]) consisting of n=120 primary tumor and n=26 metastatic samples.

### 2.2. Bioinformatic Analysis

To get a more comprehensive view, t-SNE plots and UMAPs were applied for the analyses, combined with three data transformation approaches. First, we used the unprocessed FPKM values – obtained by normalizing the mapped sequencing reads – log10 transformed values, and log10+1 transformed values. The so-called log10 transformed values, are the log10 values of the unprocessed FPKM values, where values equal to 0 are set to 0. The so-called log10+1 transformed values are the log10 values which are obtained after the unprocessed FPKM value plus 1 has been calculated.

Subsequently, results of t-SNE and UMAP dimension reduction were compared. t-SNE plots were created based on a principal component analysis with 50 components, a learning rate of 300, and a perplexity of 27. Further details on the procedure are given elsewhere [21]. UMAP plots were generated based on an adapted UMAP approach as previously described [20]. In brief, the squared pairwise Euclidean distance was used to calculate the distance between samples with a subsequent binary search for the optimal rho based on a fixed number of 15 nearest neighbors. The symmetry calculation was simplified, by dividing the sum of probabilities by 2. Furthermore, mind_dist = 0.25 was used, as well as cross-entropy as cost function with normalized Q parameter. Last, gradient descent learning was used with 2 dimensions and 50 neighbors, were applicable (NEP-WCM dataset used 25 neighbors). After generating the unbiased low-dimensional representations of the high-dimensional input (RNA-sequencing), data manual cluster interpretation was performed.

## 3. Results

### 3.1 Analysis of the PRAD-SU2C (Dream Team) Dataset

The first dataset in our analysis represented metastatic prostate carcinoma. Within t-SNE plots, up to three clusters were observable, according to applied data transformation. Unprocessed FPKM values resulted in one visible cluster in addition to the big main cluster (Figure **1** a), whereas log10 (Figure 1b) and log10+1 (Figure 1 c) approaches showed two additional smaller clusters. These clusters mainly contained bone or liver samples and were named accordingly.

**Figure 1.**
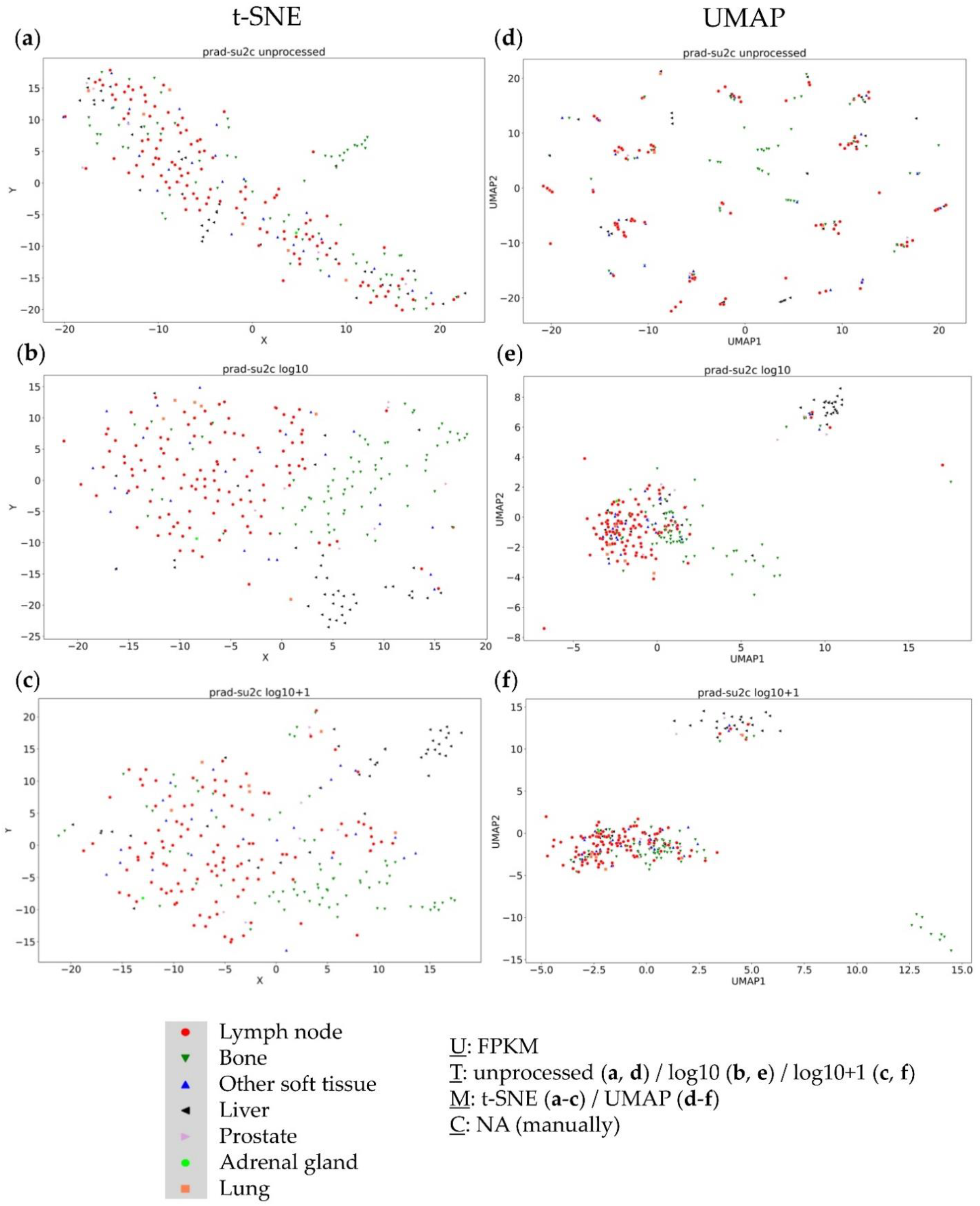
Visual clustering of the DreamTeam dataset consisting of metastatic prostate cancer (with respective resection sites) by applying different data dimension reduction methods. t-SNE plot approach for (**a**) unprocessed, (**b**) log10 transformed, and (**c**) log10+1 transformed FPKM values and UMAP approach using (**d**) unprocessed, (**e**) log10 transformed, and (**f**) log10+1 transformed FPKM values. FPKM: Fragments Per Kilobase Million; U: Unit, T: Transformation, M: data dimension reduction Method, C: Clustering method, NA: not applicable.

The UMAP approach showed similar results, with unprocessed FPKM values (Figure **1** d) not providing any clustering information, whereas log10 (Figure 1 e) and log10+1 (Figure 1 f) transformations showed three visible and distinct clusters. Again, one of these clusters completely consisted of bone samples, another mainly consisted of liver samples and the last and largest cluster consisted of all remaining samples. These results indicate that the resection site was not the main cause for clustering, instead visualization techniques (t-SNE vs. UMAP) and data transformation (unprocessed vs. log10 vs. log10+1 transformed data) heavily affected clustering results.

### 3.2 Analysis of the NEPC WCM (Neuroendocrine Prostate Cancer) Dataset

The second dataset represented neuroendocrine prostate cancer. No clusters were detectable using the t-SNE approach (Figure 2 a-c). However, the UMAP approach consistently revealed three distinct clusters (Figure 2 d-f). Of note, resulting clusters were very similar throughout all data transformations – thereby not displaying any resection site specificities.

**Figure 2.**
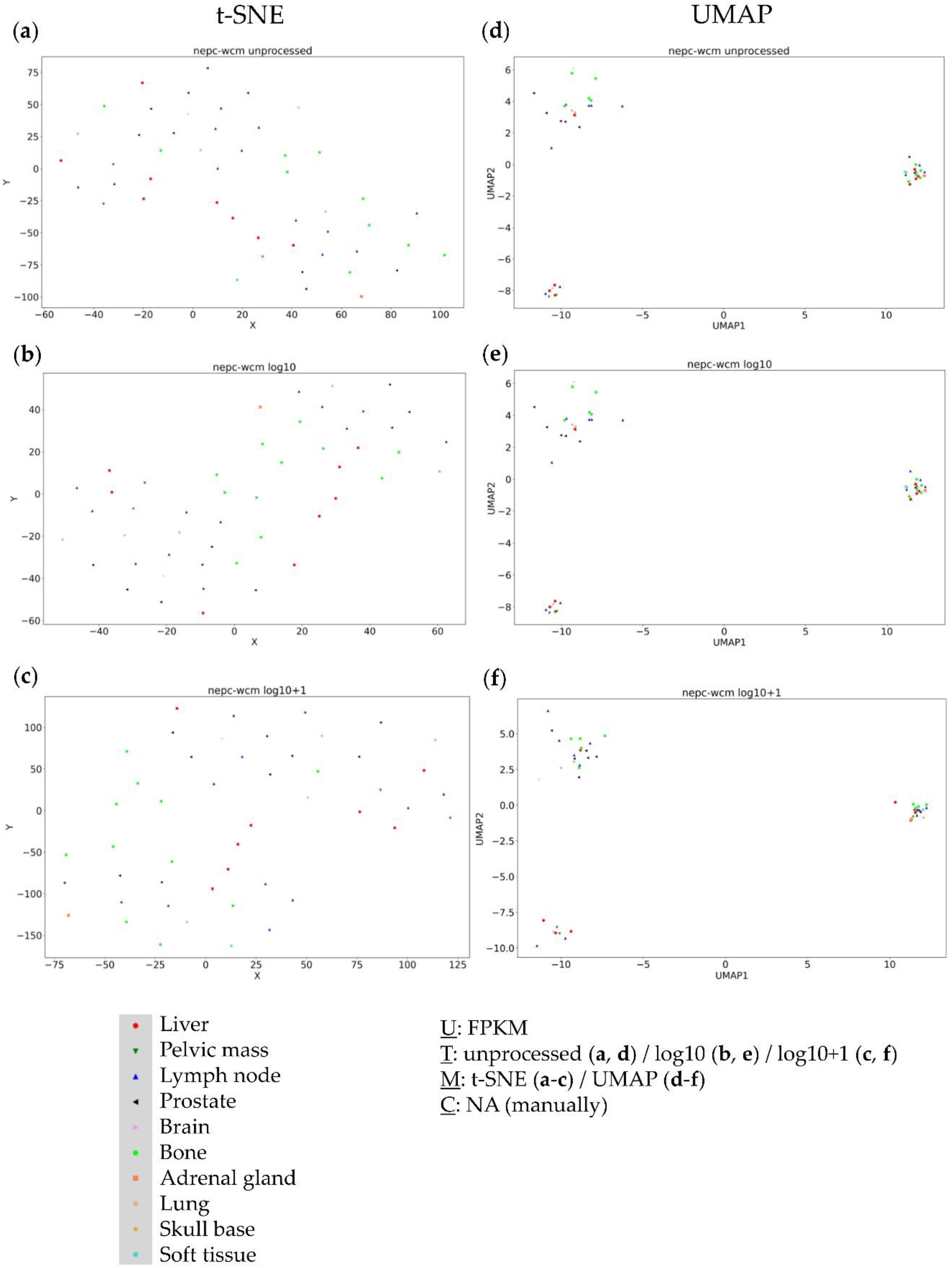
Visual clustering of the NEPC WCM dataset consisting of neuroendocrine metastatic prostate cancer (with respective resection sites) by applying different data dimension reduction methods. t-SNE plot approach for (**a**) unprocessed, (**b**) log10 transformed, and (**c**) log10+1 transformed FPKM values and UMAP approach using (**d**) unprocessed, (**e**) log10 transformed, and (**f**) log10+1 transformed FPKM values. FPKM: Fragments Per Kilobase Million; U: Unit, T: Transformation, M: data dimension reduction Method, C: Clustering method, NA: not applicable.

### 3.3 Analysis of the Metastatic Samples of TCGA-SKCM Dataset

The SKCM-TCGA dataset representing metastatic melanoma served as third dataset. Again, no clusters were detected using t-SNE plots with the different data transformations (Figure 3 a-c). Considering the UMAP approaches, unprocessed FPKM values did not provide any useful clustering information (Figure 3 d). The log10 transformed values formed one large cluster containing nearly all samples with only few outliers (Figure 3 e). Only log10+1 transformed values formed distinct clusters without site-specific agglomeration (Figure 3 f), again showing the critical impact of data transformation on cluster formation.

**Figure 3.**
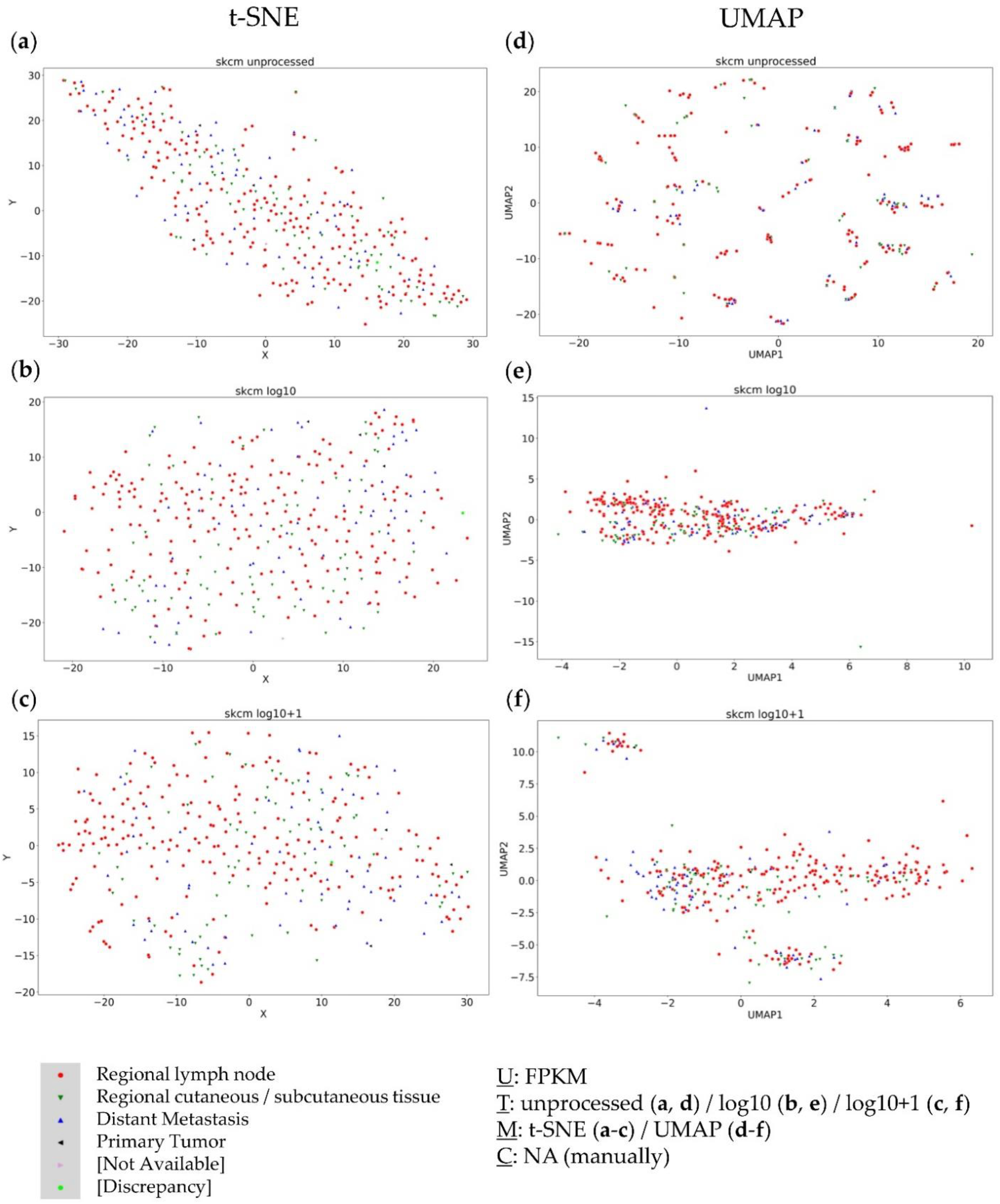
Visual clustering of the metastatic TCGA-SKCM dataset consisting of melanoma metastases (with respective resection sites) by applying different data dimension reduction methods. t-SNE plot approach for (**a**) unprocessed, (**b**) log10 transformed, and (**c**) log10+1 transformed FPKM values and UMAP approach using (**d**) unprocessed, (**e**) log10 transformed, and (**f**) log10+1 transformed FPKM values. FPKM: Fragments Per Kilobase Million; U: Unit, T: Transformation, M: data dimension reduction Method, C: Clustering method, NA: not applicable.

In summary, in none of the three datasets, a continuous dependence of the resection site could be seen. Instead, a strong dependence of the visual cluster formation on the applied method and data transformation was observed. Only one small bone cluster in the Dream Team dataset was detectable independent of data transformation and dimension reduction approaches. To validate the obtained results and to test previous observations of subgroup-dependent clustering, the KIPAN dataset – consisting of three known biologically distinct subgroups of renal cell carcinoma (RCC) – was additionally considered.

### 3.4 Further Evaluation of Cluster Formation based on Data Dimension Reduction Methods and Data Transformations

To investigate the influence of data transformation and data dimension reduction methods on forming visually distinct clusters, the TCGA data sets of the three largest RCC subgroups – clear cell (KIRC), papillary (KIRP), and chromophobe (KICH) – were combined to one data set (KIPAN). Due to the nature of the histopathologic origin of the samples in this dataset, a specific clustering could be expected.

t-SNE (Figure 4 b, c) and UMAP (Figure 4 e, f) approaches based on log10 or the log10+1 transformed data yielded a separation of samples matching the histopathologic expectation. Further, the importance and clinical relevance of subgroups identified by t-SNE (Figure 4 a) using unprocessed data for the TCGA-KIPAN dataset has already been shown [21]. Yet, the unprocessed FPKM values yielded no useful information regarding the resection site-specific agglomeration of samples in the UMAP approach (Figure 4 d).

**Figure 4.**
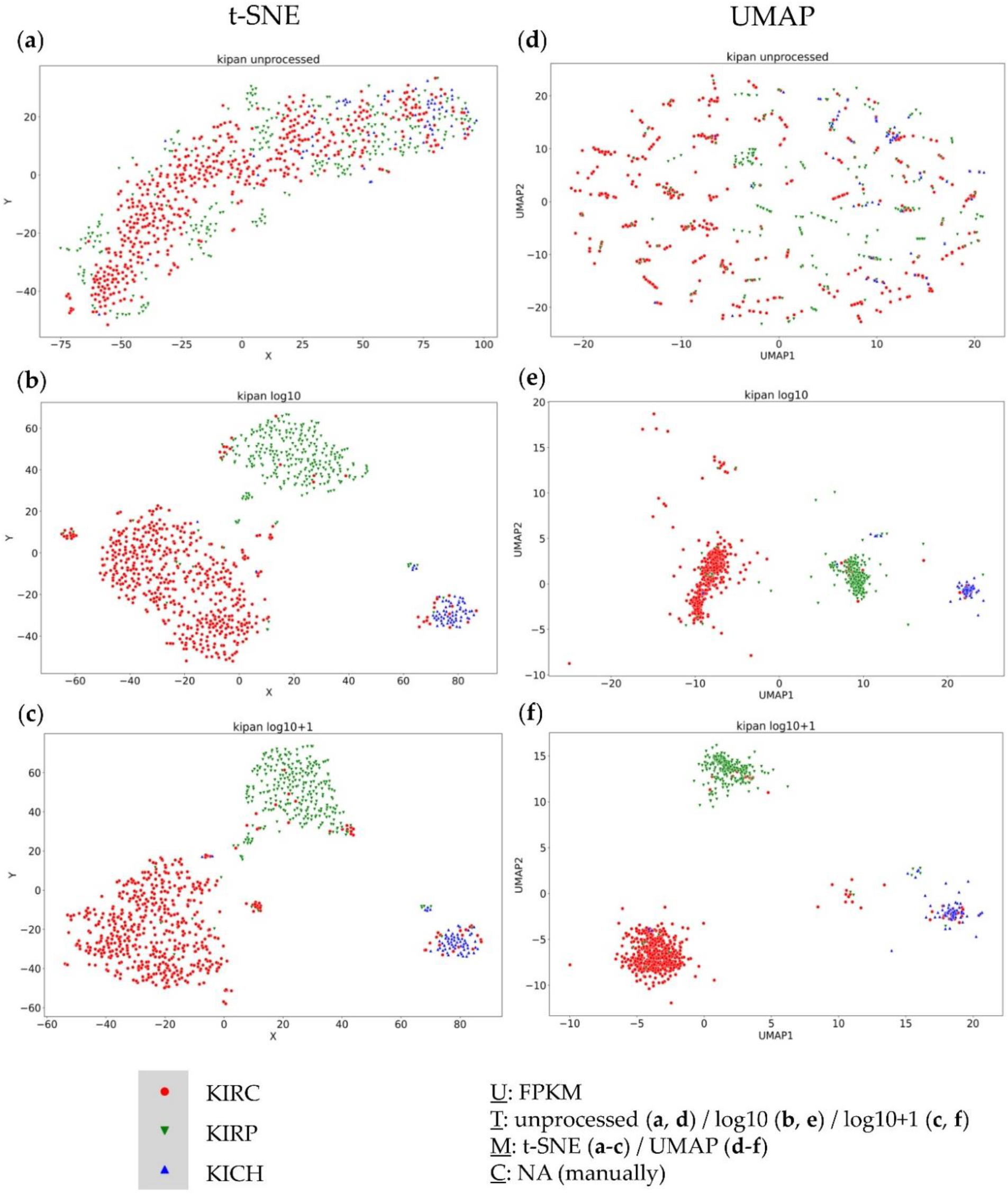
Visual clustering of the TCGA-KIPAN dataset consisting of the three major histopathologic subgroups of renal cell carcinoma (RCC) – clear cell RCC (KIRC), papillary RCC (KIRP), and chromophobe RCC (KICH) – by applying different data dimension reduction methods. t-SNE plot approach for (**a**) unprocessed, (**b**) log10 transformed, and (**c**) log10+1 transformed FPKM values and UMAP approach using (**d**) unprocessed, (**e**) log10 transformed, and (**f**) log10+1 transformed FPKM values. FPKM: Fragments Per Kilobase Million; U: Unit, T: Transformation, M: data dimension reduction Method, C: Clustering method, NA: not applicable.

Although both data transformations showed a separation based on the histopathological subgroups for both data dimension reduction methods, clusters were not exclusively subgroup-specific and displayed certain outliers.

### 3.5 Combined Analysis of Primary and Metastatic Samples of the same Entity

Based on the results of the KIPAN cohort, further analyses were performed for the complete TCGA-SKCM dataset as well – to analyse the transcriptomic relation of primary and metastatic melanoma. Interestingly, no distinct separation between metastatic and primary melanoma samples was observable (Figure 5).

**Figure 5.**
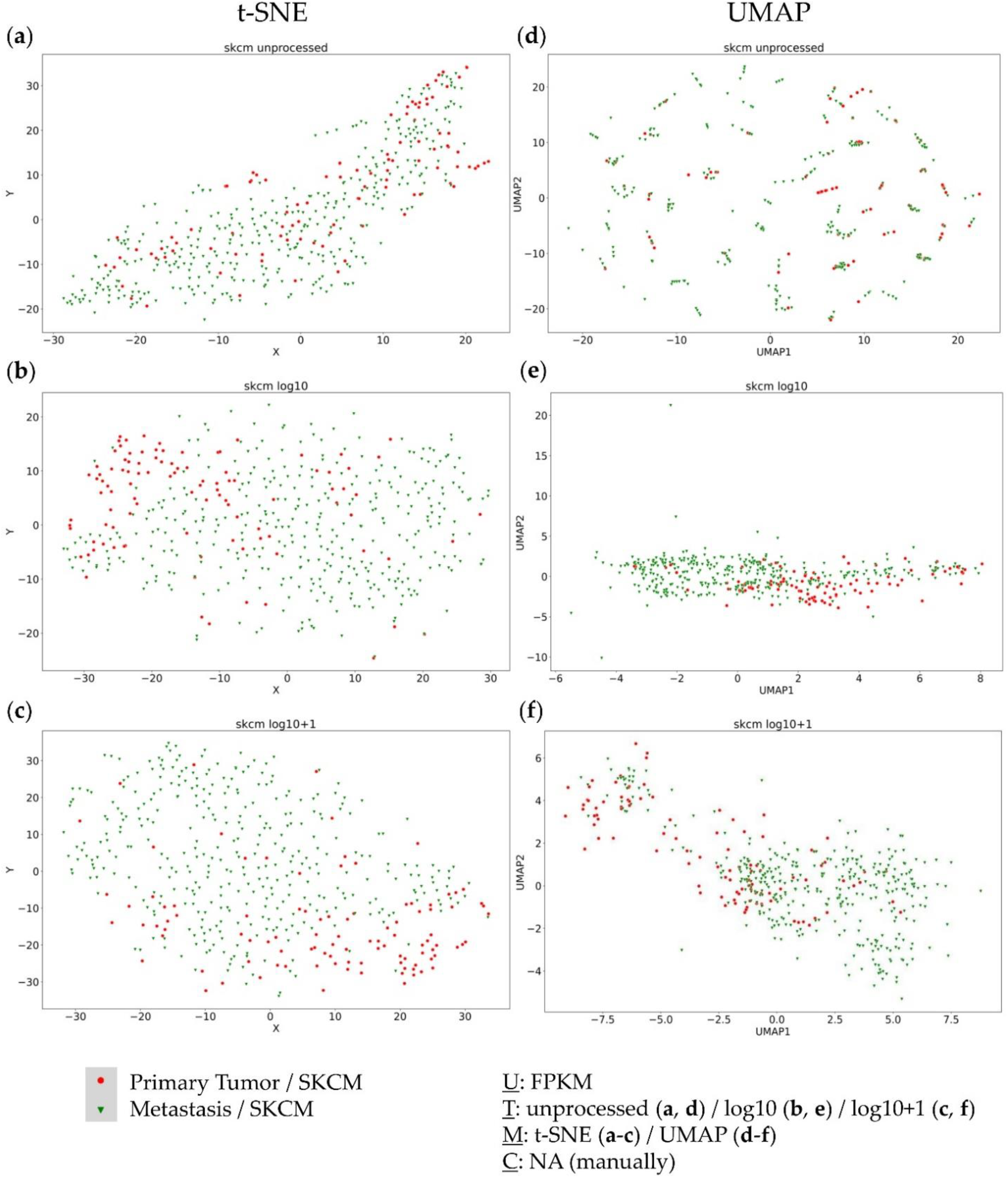
Visual clustering of the complete TCGA-SKCM dataset consisting of primary tumors (red) and metastases (green) by applying different data dimension reduction methods. t-SNE plot approach for (**a**) unprocessed, (**b**) log10 transformed, and (**c**) log10+1 transformed FPKM values and UMAP approach using (**d**) unprocessed, (**e**) log10 transformed, and (**f**) log10+1 transformed FPKM values. FPKM: Fragments Per Kilobase Million; U: Unit, T: Transformation, M: data dimension reduction Method, C: Clustering method, NA: not applicable.

Moreover, only the UMAP log10+1 transformed approach displayed two distinct clusters, each containing primary and metastatic samples (Figure 5 f). For both clusters, no complete subgroup-specific (primary tumor vs. metastasis) clustering resulted, yet a certain gradient is observable, indicating transcriptomic differences between primary and metastatic tumors, but without previous knowledge of the subgroup, no assumptions could be made in separating both groups. Moreover, some metastatic tumors seem to still harbour primary tumor transcriptomic features, whereas there are also primary tumors already harbouring metastatic features.

To further validate these results, we finally analysed the metastatic breast cancer project (MBC Project) dataset, consisting of both primary and metastatic tumors of different resection sites. Using the t-SNE approach on this dataset did not lead to cluster formation for any of the data transformations (Figure 6 a-c). Again, unprocessed FPKM values in combination with the UMAP did not provide any useful information about the dataset (Figure 6 d). Additionally, logarithmic transformations within UMAP approaches did not form any distinct clusters, either (Figure 6 e, f) – thereby confirming the findings from the TCGA-SKCM dataset.

**Figure 6.**
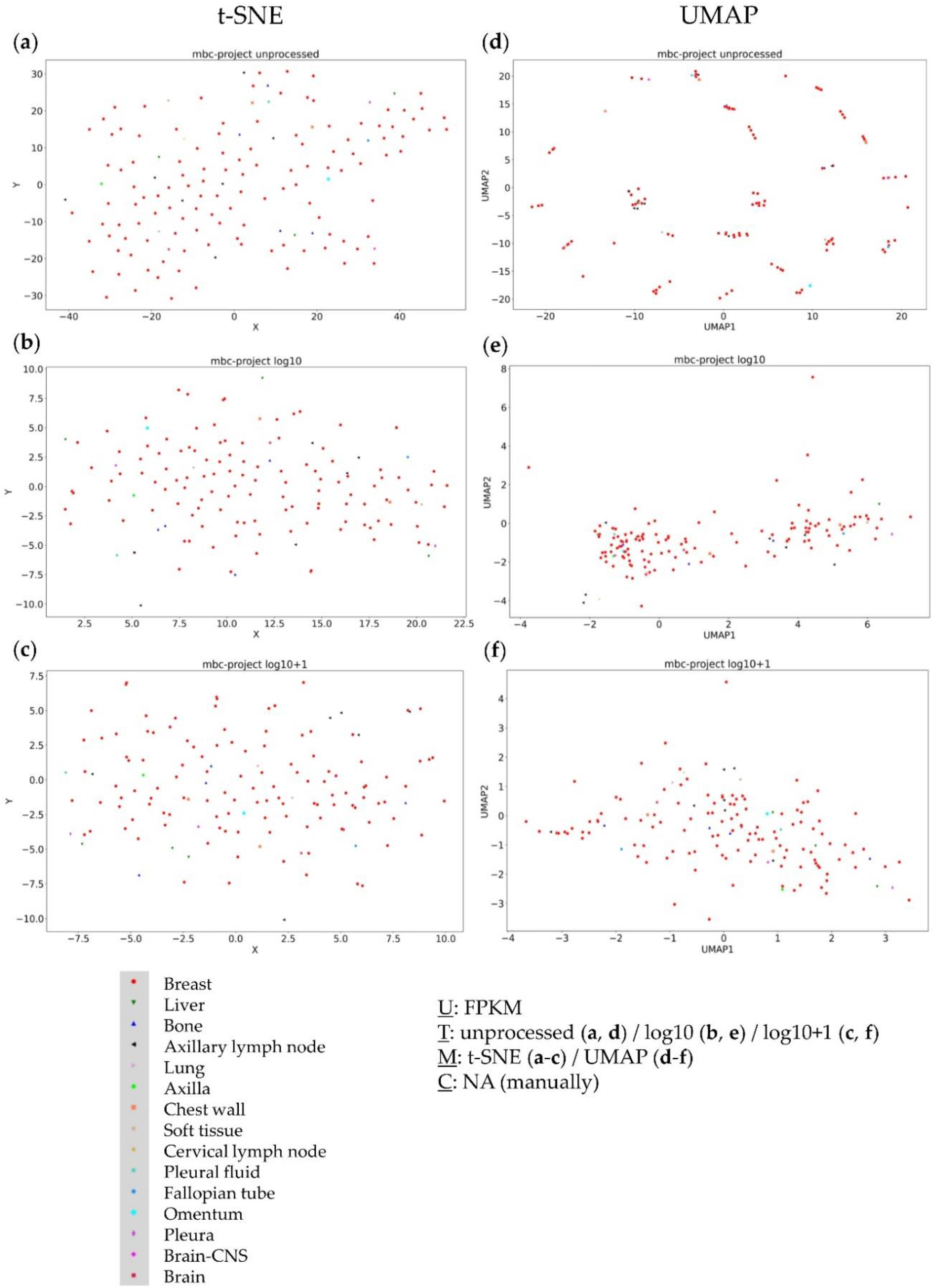
Visual clustering of the MBC Project dataset consisting of primary and metastatic breast cancer (with respective resection sites) by applying different data dimension reduction methods. t-SNE plot approach for (**a**) unprocessed, (**b**) log10 transformed, and (**c**) log10+1 transformed FPKM values and UMAP approach using (**d**) unprocessed, (**e**) log10 transformed, and (**f**) log10+1 transformed FPKM values. FPKM: Fragments Per Kilobase Million; U: Unit, T: Transformation, M: data dimension reduction Method, C: Clustering method, NA: not applicable.

## 4. Discussion

### 4.1 The Impact of Data Transformation on Cluster Formation within Data Dimension Reduction

In this work, we were looking for transcriptomic similarities and differences of metastasis representing different resection sites. It has already been shown in several studies that there is no clustering of samples depending on the underlying resection site of the metastasis [31]. Nevertheless, within these studies, clustering was frequently observed [32]. These clusters were often attributed to biologically distinct subgroups in one entity – also stating preferred metastasis sites for different subgroups [1]. Additionally, there are studies showing transcriptional differences between two different resection sites [10]. Due to this, we compared the clustering results of three different datasets. Since previous analyses did not specifically investigate the transcriptomic dependency of the resection site, our approach considered not only different unbiased data dimension reduction methods – subsequently used for visual clustering – but also different data transformations. It could be observed that especially log10+1 transformed data frequently resulted in a clearer and more distinct cluster formation when analysed with UMAP. In line with this observation, UMAP analysis of the TCGA-KIPAN dataset showed a cluster dependency mainly based on underlying RCC histopathology. However, histopathological clustering was evident in the UMAP log10+1 and in the UMAP log10 data transformations as well as in the results of the corresponding t-SNE approaches.

As already shown in a previous publication, obtained clusters by using unprocessed FPKM values in a t-SNE approach yielded prognostically relevant clusters with biologically distinct characteristics for RCC [21]. These findings were also in line with previous literature [33]. Additionally, using UMAP data dimension reduction with logarithmically transformed data of the TCGA-ACC (adrenocortical carcinoma) dataset revealed two clusters closely matching the already known ACC subgroups [20]. This suggests that histopathological and cancer subgroup-specific differences can be represented with a UMAP log10+1 approach, even though clusters seen within TCGA-KIPAN analysis were not completely subgroup-specific, also observed in t-SNE plot using unprocessed data. Since t-SNE and UMAP show biologically meaningful clustering results – known histopathological or cancer entity subgroups – based on different data transformations, both data dimension reduction methods are useable and valid, depending on the underlying biological question.

Another remarkable element is the bone cluster identified within the Dream Team dataset. This cluster appears – with minor changes and depending on the area considered – in each of our analyses, regardless of the data transformation and the data dimension reduction method. Considering previous results, we conclude that all present methods have their justification and can be used depending on the research question. For example, the UMAP log10+1 approach is suitable for bulk RNA sequencing to identify subgroups within specific entities. However, clusters based on different histopathological tissues, for example, and thus generally showing a quite different transcriptome, can also be seen in the unprocessed data, where t-SNE plot seems to be more suitable for bulk RNA sequencing than UMAP, which in turn does not seem to be suitable for unprocessed data of bulk RNA-sequencing in general.

When looking at the number of clusters previously identified for the analysed datasets within this work, it becomes apparent that log10+1 approaches were mostly in line with previously shown results. For the TCGA-SKCM dataset, three clusters were identified in in the first publication of this dataset [27]. The initial description of the NEP-WCM dataset showed three (based on the main branches of the dendrogram) different main clusters based on unsupervised clustering, overlapping with our UMAP results. Additionally, a smaller neuroendocrine subgroup inside the Dream Team dataset was described, which might be one of the shown clusters in our approach. This further proves the clusters found by unsupervised clustering in the Dream Team original publication, stating the independence of the metastasis site [25], and confirming the different molecular phenotype of neuroendocrine prostate cancers [34]. Regarding breast cancer metastases, our results confirm previous findings showing the cluster dependency on biological subgroups rather than on resection site [31].

Looking more closely at the differences in the resulting clusters between the different data transformations of the individual data dimension reduction methods, changes are similar to the ones caused by parameters such as the number of neighbors. Considering very recent research, we believe that the data transformation used is just an equally important factor to consider in the initialization of the data [23], as respective kernel transformations [24].

### 4.2 Primary Tumors and Metastases of the same Entity share Common Transcriptomic Features

Our findings that primary and metastatic tumors share common transcriptomic features and are inseparable when analysed with data dimension reduction methods appear to match with previous research. In metastatic pancreatic adenocarcinoma, a distinction between primary tumor and metastatic tumor cells was not possible using single cell RNA-sequencing [35,36]. This could also be seen in breast cancer single cell RNA-sequencing comparing lymph node metastasis with primary tumors [37], which is in line with our findings regarding the MBC project dataset, not forming visual clusters in any considered approach. The presented results support the linear progression model to some extent, at least for the transcriptomic differences between metastasis and primary tumors, indicating the need for further research to combine genomic alterations with transcriptomic features to clarify the (clonal) evolution of metastasis. In conclusion, our results suggest that there is no general transcriptomic dependency on resection site for metastasis of the same primary tumor and that obtained clusters can be mostly attributed to existing subgroups. The genetic diversity, using bulk sequencing and analytical deconvolution is a major hallmark of cancer in general. Pre-metastatic and pre-treatment diversity can help to predict the clinical and evolutionary outcome of the disease. Nonetheless, the regulatory wiring that underpins the metastatic process is likely to dynamically change across the transcriptional landscape.

### 4.3 Addressing Pitfalls in Visual Clustering

To address these challenges, we propose an additional standard-legend for visual clustering approaches based on data dimension reduction methods and machine learning, as represented by the UTMC legend in all figures of this work. The information required by this additional information includes the unit (U) (such as FPKM, TPM, RPKM or read counts), data transformations (T), represented visualization or data dimension reduction method (M) and, if applicable, the applied cluster identification algorithm (C). This enables reproducibility of figures and analyses and makes visual clustering approaches much more transparent.

Taken together, our work further extends the knowledge of tumor heterogeneity in different biological context [38], by providing sufficient evidence for the linear progression model of metastasis, since no dependency of clusters based on resection site was observable in any of the three considered datasets. The applied transformation tended to have the biggest impact on clustering results, and thus needs more in-depth analysis. Nevertheless, our results cannot identify a favourite approach, as all of them appear to properly address different questions. Transformed data, independent of the data dimension reduction method, tend to visualize subgroups very specifically, whereas using unprocessed data in t-SNE seems to be closer to the biological nature of samples, demonstrating the need for further research in this area.

## 5. Conclusions

Using two different data dimension reduction methods, we showed that there was no visual association between resection site and the transcriptome for three considered metastatic datasets. Instead, there was a significant dependence of clustering according to data transformation and the data dimension reduction method applied. Additionally, the analysis of primary and metastatic samples of specific entities did not show distinct clusters or visible differences. Combining recent works and the results of our study, visual clustering seems highly vulnerable towards data and parameter alterations. To avoid pitfalls in analyzing visual clustering and to enhance reproducibility, we recommend extending the standardized nomenclature, e. g. by adding the UTMC-legend introduced in this manuscript.

## Author Contributions

Conceptualization, A.M., A.K., A.G.S., and M.Kr.; methodology, A.M. and P.K.; software, A.M., validation, A.K., M.Kr., and M.Kn.; formal analysis, A.M. and P.K.; investigation, A.K., M.Kr., and M.Kn.; resources, A.G.S., A.K. and M.Kr.; data curation, A.M. and P.K.; writing—original draft preparation, A.M., P.K., M.Kr., M.Kn., A.G.S., A.A., AK; writing— review and editing, A.M., P.K., M.Kr., A.G.S., A.A., A.K.; and M.Kn.; visualization, A.M., A.K., A.G.S., and M.Kr..; supervision, M.Kr. and P.K.; project administration, A.K., M.Kr., P.K., A.G.S. and A.K.; funding acquisition, A.G.S. and M.Kr. All authors have read and agreed to the published version of the manuscript.

## Funding

This project was supported in part by the Apulian Regional Project Medicina di Precisione to A.G.S. Moreover, MK was funded by a personal grant from Else-Kröner-Foundation (Else Kröner Integrative Clinician Scientist College for Translational Immunology, University Hospital Würzburg, Germany). The funding source has not been involved in study design, collection, analysis and interpretation of data, writing of the report or in the decision to submit the article for publication.

## Institutional Review Board Statement

Ethical review and approval were waived for this study due to the in-silico and re-analysis nature of this study; therefore, no primary material was used.

## Informed Consent Statement

Patient consent was waived due to the in-silico and re-analysis nature of this study; therefore, no primary material was used.

## Data Availability Statement

All datasets in this study are publicly available. Datasets were either accessed via GDC-portal (https://portal.gdc.cancer.gov/projects) or cBioPortal (https://www.cbioportal.org/) [28,29].

Jupyter Notebook containing the source code of the altered UMAP approach can be requested from the corresponding author A.M.

## Acknowledgments

The results shown here are in part based upon data generated by the TCGA Research Network: https://www.cancer.gov/tcga.

The results demonstrated here include the use of data from The Metastatic Breast Cancer Project (https://www.mbcproject.org/), a project of Count Me In (https://joincountmein.org/).

## Conflicts of Interest

The authors declare no conflict of interest. The funders had no role in the design of the study; in the collection, analyses, or interpretation of data; in the writing of the manuscript, or in the decision to publish the results.

## Notes

### Competing Interest Statement

The authors have declared no competing interest.

